# Lost to science for 126 years: Indigenous Knowledge and Camera Trapping Document the Critically Endangered Black-Naped Pheasant-Pigeon *Otidiphaps insularis*

**DOI:** 10.1101/2024.12.17.628566

**Authors:** Jason J. Gregg, Jordan Boersma, Doka Nason, Elimo Malesa, Cosmo Le Breton, Serena Ketaloya, Bulisa Iova, John C. Mittermeier

## Abstract

We combined questionnaires collecting local ecological knowledge with passive and directed camera trapping and bird surveys to scientifically document the Black-Naped Pheasant-Pigeon *Otidiphaps insularis* for the first time since 1896. This species is endemic to Fergusson Island, in eastern Papua New Guinea, and is New Guinea’s most endangered terrestrial bird species. Indigenous subsistence-based hunters who participated in this research as informants had high levels of local bird knowledge and helped direct camera trapping for one of two successful camera trap captures. Local participants also shared traditional ecological knowledge about the Black-Naped Pheasant-Pigeon, including legends, chants, and a local name for this taxon, Auwo. The scarcity of observations over the course of a month-long expedition, its obscurity to the majority of local respondents interviewed, and prevalent threats from habitat destruction and introduced alien species further support its current Red List status (CR). Our results provide a case study for leveraging local ecological knowledge in combination with standardized survey techniques and may be an especially effective strategy for determining the status of other undocumented or data-deficient species in the southwest Pacific.

## Introduction

For more than twelve decades *Otidiphaps insularis* (Black-Naped Pheasant-Pigeon) has been lost to Western science. Known only from Fergusson Island, a satellite island of eastern New Guinea, the syntype specimens were collected by Andrew Goldie in 1882 (Godman and Salvin 1883, (NHMUK 1889.2.12.119 and NHMUK 1889.2.12.484). It was again collected by Albert Meek in 1896 (AMNH 616494) after which it remained completely absent from the scientific record (Gregg et al. 2020, Kirwan and van Grouw 2023). The total population of *O. insularis* is estimated to be fewer than 249 individuals and thought to be declining due to habitat loss (IUCN 2023). Given its undocumented status and risk of extinction, Gregg et al. (2020) previously recommended an urgent, island-wide survey to determine the status of the species. In 2021, *O. insularis* was subsequently uplisted by the International Union for the Conservation of Nature Red List (IUCN), becoming New Guinea’s only critically endangered (CR) terrestrial bird species (IUCN 2023). Given its unique evolutionary lineage, the taxon was ranked in the top 3% of all extant bird species when evaluated for evolutionarily distinctness and globally endangered (EDGE) criteria (McClure et al. 2023), and with no confirmed documentation in more than 10 years, the species was also identified as “lost” by the Search for Lost Birds initiative (Rutt et al. 2024).

Fergusson Island (hereafter Fergusson) is the largest island in the D’Entrecasteaux Archipelago (1,437 km^2^; max. elevation 2,073 m), in Milne Bay Province, Papua New Guinea (hereafter PNG). The islands of southeastern New Guinea are more poorly surveyed relative to other areas of the region (Frith and Beehler 1998), and ornithological surveys in the D’Entrecasteaux Islands over the last two decades have documented dozens of new avian species records for the archipelago, including a novel *Zosterops* taxon. (Pratt 2014, Gregg et al. 2020, Boersma et al. 2024, Pratt unpublished).

*Otidiphaps*, order Columbiformes, is a genus endemic to New Guinea. The four recognized taxa are large-bodied and primarily terrestrial, with long legs and a laterally folded tail of 20–22 feathers, unlike any other Columbid (Kirwan et al., 2023). The genus also claims distinct internal anatomy, including a narrow, elongated sternum and a small furcula relative to other birds (Glenny and Amadon 1955, Gibbs et al. 2001). Originally described as four distinct species, *Otidiphaps* populations have been treated as four subspecies for a century (Peters 1934, Mayr 1941, Pratt and Beehler 2014). More recently, del Hoyo and Collar (2014) again recognized these populations as distinct species using the Tobias et al. (2010) criteria, a designation used by Birdlife International and the IUCN (del Hoyo and Collar 2014, Tobias et al. 2010, Kirwan and van Grouw 2023). In the absence of a molecular phylogeny, species limits and taxonomy remain uncertain, and some authorities continue to treat all *Otidiphaps* taxa as belonging to a single species, *O. nobilis* (e.g., Clements et al. 2023). For the purpose of this publication, we follow del Hoyo and Collar (2014) and treat named *Otidiphaps* taxa as full species.

There is little information published about *Otidiphaps* in the wild, especially the two taxa existing on New Guinean satellite islands, *auruensis* and *insularis*. Restricted to Fergusson, *insularis* is the only member of the genus to occur off the New Guinea continental shelf (Bintanja et al. 2005), and is known from just three specimens. In comparison with its congeners, *O. insularis* has no backwards facing crest, no obvious contrasting coloration on its nape, and less coloration on its underbody. Contrasting with these duller features, its cadmium orange wings are noticeably brighter than congeners. Godman and Salvin (1883) provide the sole information about *O. insularis* from life, including brief notes on where it was collected (on an “exceedingly rough range of mountains” above 600 m ASL) and its vocalizations (described as “a sort of *ké-o*, the ‘o’ being prolonged” by A. Goldie during a collecting expedition). Thus, gathering additional information on the status and population size of *O. insularis* is vital.

Local ecological knowledge (LEK) is a utilized, site-specific knowledge based on interactions with local environments which is held by Indigenous and non-Indigenous peoples (Joa et al. 2018). LEK is generally proposed as epistemologically distinct from scientific knowledge, but can contain information useful for scientific or management applications. In partial contrast, traditional ecological knowledge (TEK) is defined as a complex of local environmental knowledge, beliefs, values, traditions, and practices (Berkes et al. 2000). A growing body of research has incorporated LEK for collecting data on rare, critically endangered, and/or cryptic species using an interview or questionnaire-based approach. LEK has been fundamental to ornithological research in New Guinea (Joseph et al. 2019), and along with strong customary land ownership and traditional land management systems, local people continue to practice subsistence-based activities upon which much of LEK and TEK are thought to be based.

Hunters in PNG are recognized as being highly knowledgeable about animal species, which they harvest for consumption and traditional uses in many parts of the country (Mack and West 2005). As one of Earth’s great epicenters of avian diversity, yet scarce of large mammals, birds are especially important to the people of New Guinea, and are regularly eaten and used for cultural purposes (Mack and Wright 1998). Research by Kik et al. (2023) has shown hunting skills are correlated with ability to name bird species. In PNG, hunters are therefore strong candidates for being key informants in research focused on terrestrial vertebrate species. Gregg et al. (2020) previously reported continued subsistence hunting practices on Fergusson, and the large size and natural history of *O. insularis* would make it a likely candidate for a game species. During informal questioning on Fergusson in 2019, (Gregg et al. (2020), reported that hunters recognized and named *O. insularis*. Accounts from biologists and anecdotal discussions with hunters near Alotau (Milne Bay), Port Moresby (Central) and other parts of New Guinea indicate *Otidiphaps* taxa are hunted on mainland New Guinea for food (I. Voxvold pers. comm., D. Wakra pers. comm.) Thus, we hypothesized that if *O. insularis* persisted on Fergusson, local people and especially hunters, would hold observational knowledge useful for documenting the species.

To determine the status of *O. insularis*, we conducted a month-long, multi-method survey on east Fergusson that included questionnaires and interviews with indigenous and local peoples, camera trapping and bird surveys dedicated to documenting this Critically Endangered species.

## Methods

### Hunter interviews

To investigate the status of *O. insularis*, and to understand local knowledge of birds and hunting practices generally, we administered questionnaires to hunters and conducted semi-directed interviews with other informants in east Fergusson. Questionnaires were conducted in proximity to (or in) mountainous areas identified as *O. insularis* habitat by descriptions of where the syntype specimens were collected and information on its congeners (Salvin and Godman 1883, Mayr and Rand 1937, Freeman et al. 2013). All surveys occurred in east Fergusson on and around the Mt. Kilkerran Massif, the island’s tallest mountain range.

Like other epistemologies, LEK is considered heterogenous among community members, and research methods dependent on informants generally target a subset of people suspected or known to hold information of interest, an example of purposive sampling (Tongco, 2007). We identified participants who self-identified as hunters, and/or were recommended to us as hunters by community members, and study participants were selected opportunistically upon the arrival of our research team in villages. Questionnaires and semi-directed follow-up interviews were conducted by JB and EM and interpreted to English by EM, a resident of east Fergusson, from the following languages and named dialects: Dobu, Basima, Galea, Bosalewa, Salakadi, as well as the lingua franca and national language of PNG, Tok Pisin. We did not record respondents’ names or contact information to ensure data remained completely anonymous.

The survey began with a brief declaration of intent to learn about hunters’ local bird knowledge, and free, prior, and informed consent was given by each participant before beginning the questionnaire. Part one of the questionnaire included demographic and hunting behavior questions. Part two of the questionnaire used visual cues to measure bird identification accuracy, assess which bird species were hunted and why. (Figure 2, A). Part three of the questionnaire asked natural history information about species identified from visual stimuli.

Previous studies focused on LEK sometimes test participants for identification accuracy to screen informants who are knowledgeable about a species of interest (e.g. Jain et al. 2024). Similarly, our questionnaire was designed to simultaneously test whether participants were able to differentiate species present on Fergusson from morphologically distinct species absent from the island and whether they were able to identify *O. insularis* from a lineup of other bird species.

To begin part two of the questionnaire, visual cues in the form of twenty-five 10 x 10 cm laminated cards showing high-quality illustrations of birds were placed randomly before each participant (Birds of the World, 2022). Cards showed species present on Fergusson, morphologically distinct species absent from Fergusson, deemed error cards, and a third category encompassing species whose distribution on Fergusson was unknown as well as species deemed difficult to identify due to having lookalikes. (For full species list and their categories, see supplementary material). Participants were asked to identify species they see in their local area, and subsequently to select the species they hunt, ranking the order of their relative importance for hunting purposes. Both selections were photographed to record results.

We then played the vocalizations of four vocal, widespread breeding bird species common in the lowlands and hill forests of Fergusson: *Megapodius reinwardt, Ducula spilorrhoa, Paradisaea decora*, and *Erythropitta macklotii*, and either or both of two *Otidiphaps* taxa (*nobilis* and *cervicalis)* from recordings sourced from Woxvold and Bishop (2020). Species were played in random order, and participants could request audio playback be repeated as many times as needed. After each playback, participants were asked to identify each species by their local name, which were verified by EM. *Otidiphaps* recordings used during playback differed in being either one or two-noted calls, though we did not track which of these two recordings was played during each interview.

Lastly, we asked participants natural history information about bird species of interest, including *O. insularis*. Initially, natural history questions were conceived as a secondary test to gauge participants’ accuracy of LEK, however in practice, we did not follow this protocol and did not collect substantial information about other bird species due to time constraints. We do not consider this portion of the questionnaire further. Instead, we used semi-structured followup interviews with participants who had identified *O. insularis* to ask about natural history information and whether they had encountered this species recently.

Bird identification accuracy was calculated as the proportion of error cards chosen out of a total of seven possible error cards. Playback identification accuracy was calculated from the proportion of vocalizations identified out of four species and did not include identification of *Otidiphaps*. These scores were combined into an aggregated bird knowledge score out of 100% used compare bird knowledge across study participants. Participants who had both high bird identification accuracy and successfully identified *O. insularis* from the lineup of other species were considered the best sources of information about the status of *O. insularis*, and qualified as candidates for helping direct targeted camera trapping and bird surveys.

A recognized phenomenon in sociological research is that participants aware of an outcome of interest may be inclined to give socially desirable responses, known as social desirability bias (Grimm, 2010). To reduce bias for false-positive identifications of *O. insularis*, we included an element of deception in the design of our questionnaire by not emphasizing our target species relative to other bird species during identification exercises. Once questionnaires were completed in each community, we used a different strategy, revealing the search for *O. insularis* as our primary research goal. Additionally, we led outreach at the village and community level to raise awareness about resource management and recruit additional informants who may be knowledgeable about *O. insularis* who did not participate in the questionnaire. To prevent the news of our primary research goal from reaching possible questionnaire participants in other villages, we maintained a rapid schedule, visiting new villages every 4-5 days. However, we found that knowledge about our target species spread rapidly, and several hunters said they knew about our primary research goal in advance of participating in the questionnaire.

### Camera traps

We deployed two models of automated camera traps at 30 non-random, unbaited locations on the Mt. Kilkerran Massif (Reconyx, Model: Hyperfire H2X, Holmen, WI, USA and Browning, Model: Strike Force Pro BTC-5DCL, Prometheus Group, Birmingham, AL, USA). Cameras were placed vertically, ∼10 cm above ground, and mounted on white, ∼1-meter-long PVC pipes impaled into the ground. The area surrounding the camera’s immediate field of view was cleared of obstructive vegetation to prevent false triggering. Cameras were placed to capture favorable microhabitat for terrestrial, frugivorous vertebrates, including game trails, funnels, fruit falls, and areas of soil disturbance attributed to foraging. Camera settings varied between models; however, both models were set to high sensitivity and rapidfire image capture, capturing either up to 5 images (Browning) or exactly 5 images (Reconyx) with no delay. A subset of Reconyx cameras simultaneously captured video upon being triggered. All images were marked with a date/time stamp.

Twelve cameras were operated along an elevational gradient between 364 – 968 m in suspected *O. insularis* habitat, with >200 m between stations. Additionally, two groups of four cameras each were operated at locations designated as harboring *O. insularis* by participants who had accurately identified *O. insularis* during questionnaires and subsequent semi-directed interviews (Figure 2). In these areas, we placed cameras according to hunters’ guidance in locations where they reported encountering *O. insularis* within the previous calendar year. Camera trap photos were managed in R (camtrapR, Niedballa et al. 2016). Camera operation times were calculated based on deployment and retrieval dates and times. Consecutive captures of the same species were considered independent captures if they were >5 min apart. All birds and mammals were identified to species or highest taxonomic level possible using commonly available field guides and GBIF (Flannery 1995, Pratt and Beehler 2014, Gregory 2017, GBIF 2024).

### Bird surveys

To survey for *O. insularis* and Fergusson’s avian community generally, we conducted daily, ad-hoc bird surveys, identifying all birds encountered by sight and sound. Surveys were conducted by one or multiple observers on an opportunistic basis in and outside suspected *O. insularis* habitat, incidentally during foot-based travel on trails, in villages, on roads, and in areas without trails. No systematic or quantitative surveys were attempted. Birds were photographed and recorded during surveys using handheld cameras and sound recorders and documented with camera traps. Bird observations and media vouchers are reported in a separate publication and are publicly available through eBird and the Macaulay Library (Boersma et al. 2024, eBird 2021, https://www.macaulaylibrary.org/).

## Results

Our search between 5-30^th^ September resulted in the successful documentation of *O. insularis* on two camera traps in East Fergusson. In one instance, LEK informed camera trap placement and resulted in the successful photographic documentation of a single *O. insularis* on September 28, 2022 (Figure 1). A second camera trap placed along an elevational gradient without LEK also documented a single *O. insularis* on September 22, 2022 with both photos and a video (Figure 2). Captures were separated by > 5 km, and are suspected to constitute two individuals. We did not record *O. insularis* during bird surveys (n = 70).

**Figure 1.**
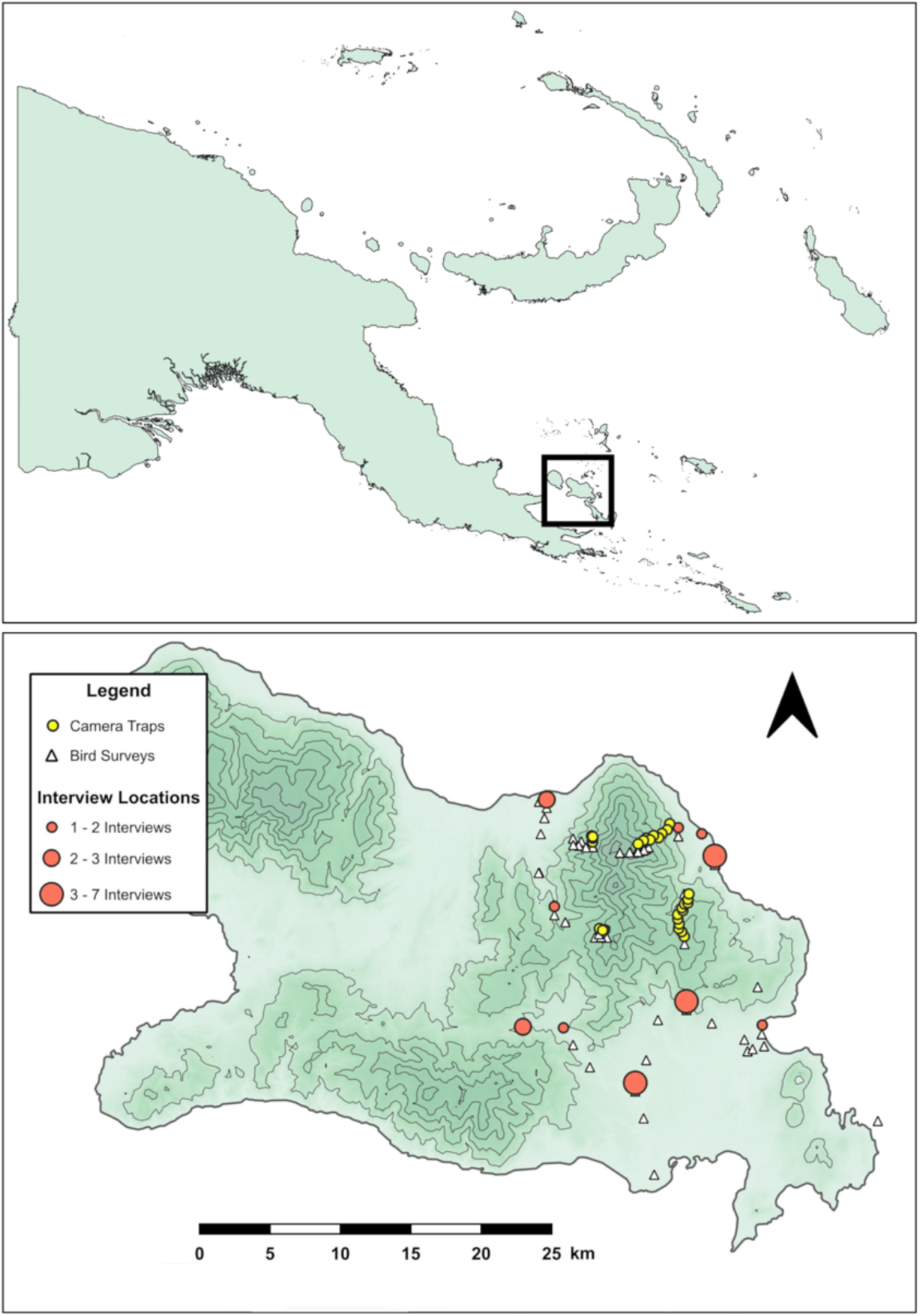
Fergusson Island. Locations of camera traps, bird surveys, and interview locations.

**Figure 2.**
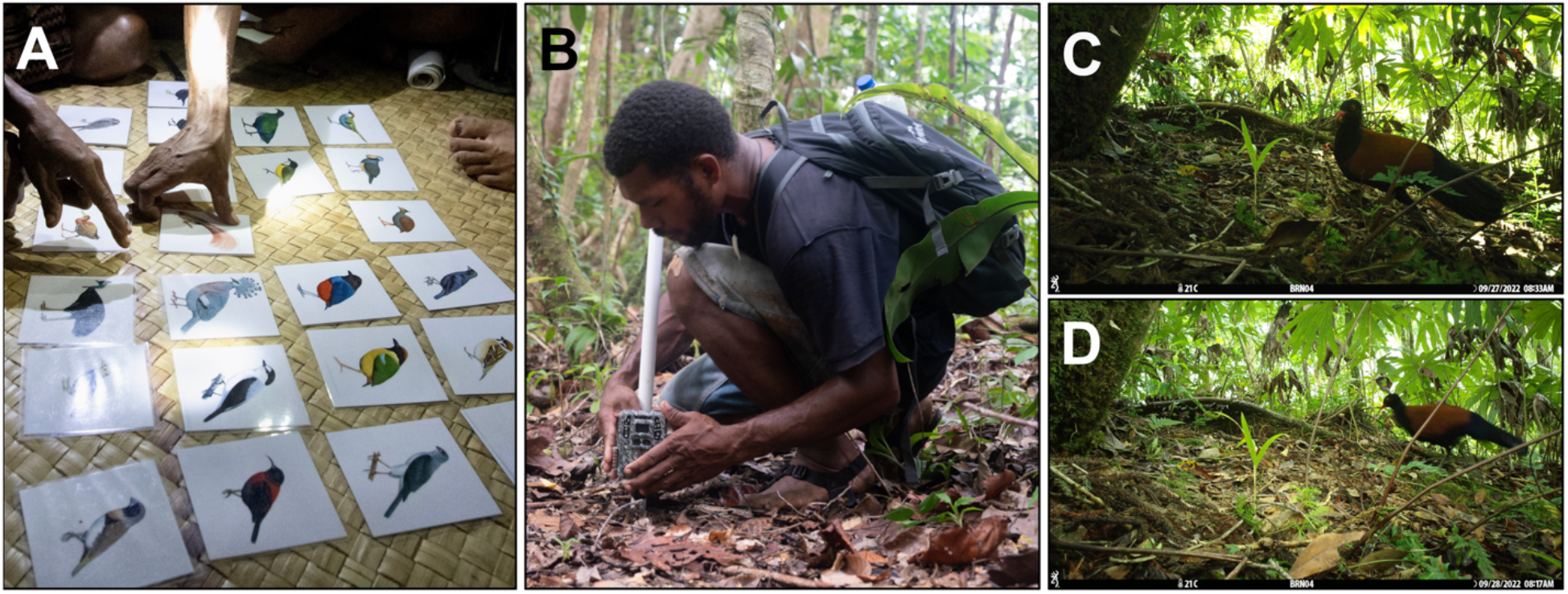
**A** Participant identifies birds he has seen on Fergusson Island, **B** Author Doka Nason sets a camera trap, **C** and **D** Two captures of the Black-Naped Pheasant-Pigeon assisted by local ecological knowledge.

**Figure 3.**
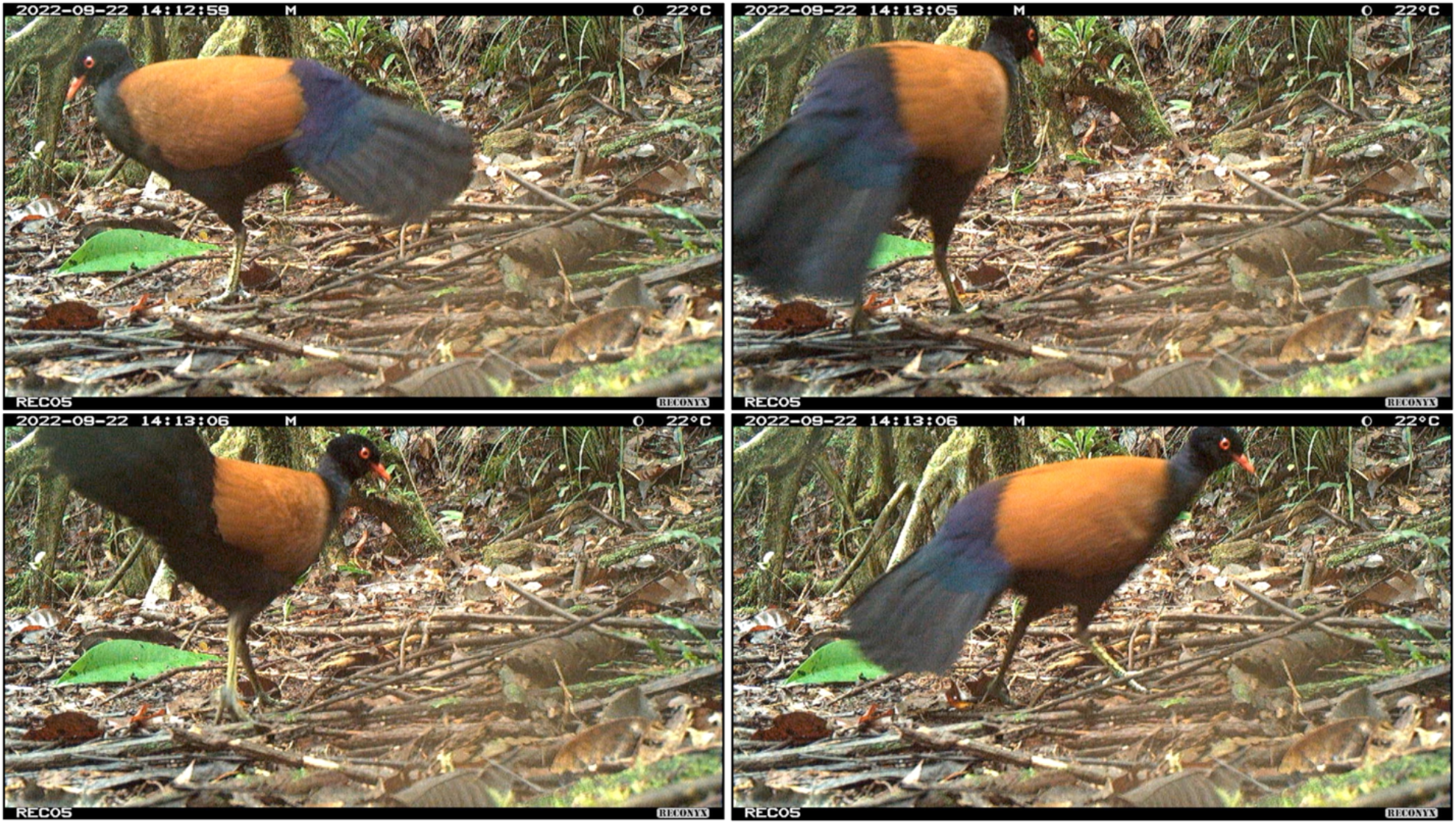
Stills from video capture of Black-Naped Pheasant-Pigeon, vigorously pumping its tail during locomotion.

### Questionnaire results

We administered questionnaires to 29 participants in ten locations surrounding the Mt. Kilkerran Massif in east Fergusson, with one to seven questionnaires completed per location (Figure 1). Questionnaire results revealed a high overall level of bird identification accuracy, a continued practice of subsistence hunting, and traditional ecological knowledge about *O. insularis*, including its role in chants, legends, and ceremonial use. Two participants reported having observed *O. insularis* within the previous calendar year.

The majority of participants (90%; n=28) reported actively hunting and 71% (n=22) responded that they hunt birds. The average frequency hunted was 9.5 days/month (range: 2-30 days/month). The average age of participants was 43 years old (range: 18-80 years old), the average age participants began hunting was 13.5 years old (range: 8-34 years old), and participants had been hunting on average for 30.5 years (range: 12-56 years).

Card identification accuracy, calculated based on the number of error cards chosen by each participant, was 84% across 29 participants, with an average of 1.1 error card chosen per participant (range = 0-5). Participants chose error cards at differing rates. Large and/or morphologically distinct species were chosen the least: *Casuarius unappendiculatus* (n=0), *Goura victoria* (n=1), and smaller and/or more colorful species were chosen the most: *Pitta versicolor* (n=12), *Ptillorrhoa leucosticte* (n=8) and *Mino dumontii* (n = 6). Identification of species during playback was 82% overall, and varied by species: *Megapodius reinwardt* and *Paradisaea decora* (identified by 93% of participants), *Ducula spilorrhoa* (75%), *Erythropitta macklotii* (61%). The mean combined identification accuracy of visual cues and audio recordings was 82% across all participants.

In contrast, 11 participants (38%) selected the *O. insularis* card, 5 (17%) identified *Otidiphaps* during audio playback of congeners, and 8 (28%) participants described the size, habitat, behavior, and field marks that matched known information about the species during semi-structured follow-up interviews. Two participants (7%) reported having observed *O. insularis* within the last calendar year.

We also report TEK about *O. insularis* that we recorded during interviews and conversations with community members. Firstly, we report the local name of *O. insularis*, Auwo, which was used by 5 participants speaking the Galea language, and which informants stated was an onomatopoeic description of the species’s vocalization. Additionally, one participant recited a chant about *O. insularis*, associating the species with love and good weather, afterwards explaining this taxon is most vocal during times of good weather. Consent to share this information was given by the participant, however a second chant including the taxon was not shared due to cultural sensitivity. This participant also recounted the collection of *O. insularis* tail feathers for use in ceremony, noting that the practice was no longer done. Another participant recounted the following legend about *O. insularis*: A village woman worked hard in her garden but was mistreated by her family, who gave her food without coconut, while they all ate food with coconut. The woman ran away from her village, travelling deep into the wilderness, where she was transformed into the Black-Naped Pheasant-Pigeon. According to the legend, the pheasant-pigeon’s call, described by this informant as human-like, is the animalized woman crying in anguish because she misses her home. Notably, both instances of TEK came from participants living on the western side of the Mt. Kilkerran Massif, the only area where we recorded encounters with *O. insularis* within the previous calendar year.

Natural history information regarding Fergusson’s avifauna was not collected consistently throughout the survey due to time limitations and our focus on our target species. We asked a subset of 16 participants about their reasons for hunting birds. All 100 % (n=16) responded that they hunted species for food, while 25%, (n=4) hunted a species due to its being associated with negative local beliefs, and 19%, (n=3) hunted a species due to its propensity for killing chickens.

Lastly, we spoke with two Fergusson residents who were identified as local historians, and who recounted an intergenerational story about meeting Andrew Goldie who was the first foreigner to collect birds on Fergusson, and whose arrival preceded most missionaries in the area. The confirmation of Goldie’s collecting efforts occurring in southeast Fergusson provided geographic confirmation regarding the general locality of where *O. insularis* was collected in 1882, information which to our knowledge is not published.

### Camera trapping results

Thirty camera traps were operated for 253 camera days producing a total of 3049 triggers, 983 captures, and 147 independent captures for a total of .6 independent captures/camera day. In total, 7 bird species (*Megapodius rheinwardti, Erythropitta macklotii, Accipiter poliocephalus, Chalcophaps stephani, Otidiphaps insularis, Colluricincla megarhyncha, Podargus ocellatus*) and 5 mammal species (*Isoodon macrourus, Rattus exulans, Rattus norvegicus, Canis familiaris, Phalanger intercastellanus)* were recorded, with 83% (n = 122) of independent captures identified to species (100% of birds were identified to species). *O. insularis* was documented on two occasions: On 22 September, a single individual was photographed on a camera trap in primary forest at 746 m ASL. This camera was placed along an elevational transect without guidance from informants, and located in primary hill forest that was visited by people hunting, collecting wood and other forest products, and travelling to villages in south and central Fergusson. The forest showed signs of light disturbance from human activities including a well-worn footpath. On 27 and 28 September, a single *O. insularis* was photographed on a camera trap set in primary forest on a narrow ridge at 971 m ASL. This camera was placed in a dense understory of ferns and other vegetation, in the exact location where an interview participant reported seeing the species. This participant had previously accurately identified *O*.*insularis* during the questionnaire, accurately described the species during followup interviews, and reported observing this taxon within the previous calendar year. This area had very few indications of human disturbance and was only occasionally visited by local landowners during extended hunting trips. These two captures were separated by > 5 km, and are suspected to constitute two individuals. We omit detailed spatial information regarding captures to reduce the risk of this taxon being targeted by the wild bird trade.

### Bird survey results

We completed 1-6 bird checklists daily, totaling 70 surveys spanning 0 - ∼1780 M ASL (Figure 1). *O. insularis* was not observed during these surveys or by other incidental observations by human observers. Ninety-three bird species were observed during surveys, including 8 species not previously documented on Fergusson, results which are reported in a separate publication (Boersma et al. 2024). Combined with surveys by Gregg et al. (2020) these efforts have resulted in 13 new island records, documented 83% (n = 100) of Fergusson’s 121 terrestrial bird species, and generated voucher media for 64% (n = 78) of these species (Boersma et. al. 2024).

## Discussion

We combined questionnaires targeting LEK with passive and directed camera trapping to document *O. insularis* for the first time since 1896, generating the first photographs and video of New Guinea’s most endangered terrestrial bird species. Understanding whether long-undocumented species persist in a rapidly changing world is vital for management interventions countering decline and extinction, and we provide a successful case study for the use of LEK for biodiversity monitoring and conservation, namely in aiding the scientific documentation of a highly cryptic bird species which remained undocumented for 126 years. By leveraging LEK as a complementary approach to standard techniques in biodiversity monitoring, research can achieve greater temporal and observational coverage than time, resource, and experience limited international research efforts can alone.

Our study adds to a body of research utilizing LEK for generating distribution data on rare, cryptic, and/or endangered species (e.g. Nash et al. 2016, Pan et al. 2018). This study also confirms the findings of research in PNG which has shown that LEK can be a scientifically accurate and operationally relevant metric for use in species conservation (Sinclair 2012). In PNG, LEK likely exceeds scientific ecological knowledge of many taxa, and we argue that connecting local and scientific knowledge networks is important for filling knowledge gaps. It may also be critical for rescuing endangered species from extinction.

We generated secondary results relevant to this poorly-known species. Our study recorded TEK and a local name for *O. insularis* in the Galea language, which we argue is valuable information for understanding the local importance and perception of this taxa, information which is likely to be relevant for the management of this species in participation with local communities. Although learning about the cultural perception of *O. insularis* on east Fergusson was not a planned research objective, we documented information about this taxon in a legend and chant in two villages. Thus, we argue that conservation designations based on evolutionary distinctiveness and/or rarity may under-rank this taxon, due to its omission of input from people who know the species best, interact with it, and may be most impacted by its decline or extinction due to importance for cultural or subsistence activities.

Notably, *O. insularis* was not detected by bird surveys in either the present study or 2019 (Gregg et al. 2020), despite the authors’ expertise with PNG and Fergusson avifauna. Working against brief and infrequent visits by foreign researchers, *Otidiphaps* are elusive, behaving like tropical forest-dwelling pheasants of SE Asia (*e*.*g. Lophura, Argusianus*). While vocalizing, the genus is readily identified, but is otherwise undercounted (B. Benz and I. Woxvold pers. comm). Vocal phenology remains unknown for *Otidiphaps*, and birds may vocalize for a short period annually, limiting planning of short-term research expeditions to coincide with when individuals are most detectable. On the other hand, LEK is defined as constituting a historically continuous body of knowledge developed over long periods of time (Joa et al 2018). Many people in Papua New Guinea engage in hunting, gardening, and other activities in the forest year-round, providing better spatio-temporal coverage of seasonal, rare, and/or cryptic species. Combined with surveys by Gregg et al. (2020), 40 total days of surveys spanning two expeditions in 2019 and 2022 covered just ∼7% of the calendar year. In contrast to these surveys, which did not document *O. insularis*, leveraging LEK with a limited array of camera traps documented this taxon over just 5.4 camera trap days.

Knowledge of *O. insularis* was heterogeneous, and the presence of *O. insularis* confirmed by camera trapping did not mean local hunters reported knowing about these species. In one instance, 7 out of 8 participants with otherwise high bird identification accuracy did not identify or report knowing this taxon, despite it being captured on a passive camera trap within ∼5 km of the village where interviews were conducted. Similarly, a study focused on LEK for recording the presence of a cryptic, terrestrial pheasant species in Asia found higher occupancy from camera traps than indicated by questionnaire results (Turvey et al. 2023). Research by Kai et al. (2014) and Turvey et al. (2018) suggest that species-specific LEK may itself track the decline and extinction of a species, a phenomenon Pyle (1978) has called “the extinction of experience”, which in turn would effectively reduce the opportunity for recording LEK and TEK during a critical phase for detection and potential conservation intervention. In our study, whether these false negatives are due to issues associated with the design of the questionnaire, the extreme elusiveness and low-density of *Otidiphaps*, or the “extinction of experience” remains unknown.

We consider the test of identification accuracy a measure of accumulated observational knowledge, which has been found to contribute scientifically accurate information elsewhere in Papua New Guinea and the Solomon Islands (Sinclaire 2012). However, using visual stimuli for measuring species identification has recognized limitations. An organism’s size, voice, behavior, and other ecological contexts are absent from two-dimensional, unscaled visual cues (as was the case in our study), and thus do not necessarily recall the first-hand observational experiences thought to underlie LEK of birds in PNG (Diamond and Bishop 1999). In a linguistic study of bird names in East New Britain Province, PNG, participants had local names of four *Accipiter* species, yet were unable to accurately identify them when shown photographs obscuring size differences (Frye et al. 2022). Scientific species may not align with local taxonomic rankings, such as ethnospecies, and local taxonomies in PNG may not identify all organisms to the species level (Frye et al. 2022). While we consider the ability of hunters to identify birds from visual and acoustic playback a useful measurement of LEK, this metric cannot be used as an index for such knowledge in general. Whether participants consider identification accuracy of birds a significant component of ecological knowledge was not investigated.

Our results support the IUCN designation of (CR) for *O. insularis*. While bird identification was high amongst interviewees, a small subset (28%) of interviewed hunters on Fergusson were able to confirm the identification of this taxon during questionnaires, with just 7% reporting observations within the past year. *O. insularis* constituted 2% of independent camera trap captures, while bird surveys over the course of two expeditions failed to document this species altogether. In contrast, another terrestrial bird present on east Fergusson, *Megapodius reinwardt*, was identified by 59 % (n=17) of hunters, made up 5% of independent camera trap captures, and was detected during 5 bird surveys.

Islands are the global epicenter of avian species loss (Fernández-Palacios et al. 2021), and large-bodied, morphologically, and functionally distinct bird species have been shown to be at highest risk of extinction (Gaston and Blackburn 1995, Hughes et al. 2022, Ali et al. 2023). We suspect logging by international corporations operating on Fergusson and the introduction of invasive alien species the most immediate conservation threats to this species. For example, timber extraction in the past two decades has resulted in logging roads constructed 3 km from the location where one individual was documented, and preliminary questioning during our interviews with hunters revealed the presence of feral cats occupying Fergusson’s forests. *O. insularis* may also be threatened by global climatic processes including long-term biogeographic kinetics. The D’Entrecasteaux Archipelago once formed a contiguous landmass called the D’Entrecasteux shelf (Diamond 1972), which subsequently fragmented with interglacial sea-level rise (Bintanja et al. 2005). *O. insularis* on Fergusson may thus represent a fragmented relict population undergoing long-term decline due to range contraction, a known fundamental driver of species extinction (Mace et al. 2008).

We recommend several areas of research to increase understanding of *O. insularis*. The current study was limited to east Fergusson, and the presence of *O. insularis* in two additional mountain ranges on the island is assumed by IUCN population estimates but unconfirmed. Considering Pleistocene land connections among the D’Entrecasteaux Islands and a highly similar avifauna with Fergusson, we recommend targeted surveys on Goodenough and Normanby Island employing our methodology, as undocumented populations may be present there. Additionally, the use of phylogenetics and modern species delimitation approaches are needed to determine the taxonomic status of *O. insularis* with greater certainty, clarifying its degree of evolutionary distinctiveness within its genus. Finally, two *Otidiphaps* taxa are widespread at zoos around the world and are able to breed in captivity (Sierra 2012). If deemed necessary to prevent extinction, we suspect a captive population of *O. insularis* could be similarly managed. However, evidence of its cultural value, as well as its suspected unique functional role amongst a depauperate island fauna strongly support *in situ* conservation. Fundamentally, any research and management activities must be conducted with the permission and will of customary landowners on Fergusson, and our study presents one avenue for working productively with local people. We hope the scientific documentation of *O. insularis* informs an effective long-term conservation strategy for this species that integrates local and indigenous knowledge and supports community capacity building, and propose our methodology as an effective approach for the search and conservation of elusive and cryptic island-endemics across the Pacific.

## Acknowledgments

We thank the residents of Fergusson Island who shared their local knowledge and hospitality with us and granted access to their land to conduct our research. Our work was possible thanks to approval from the Papua New Guinea National Research Institute and the Milne Bay Provincial government. The project was funded by the RIDGES foundation with support from American Bird Conservancy. We are also grateful to D. Mitchell of Eco Custodian Advocates, T. Pratt, and G. Dutson for sharing their expertise on D’Entrecasteaux birds and logistics. We also thank Johnila Nabai of East New Britain, who translated the abstract into Tok Pisin.

## Supplementary Material

### Supplementary Material 1

Interview questionnaire, part 1.

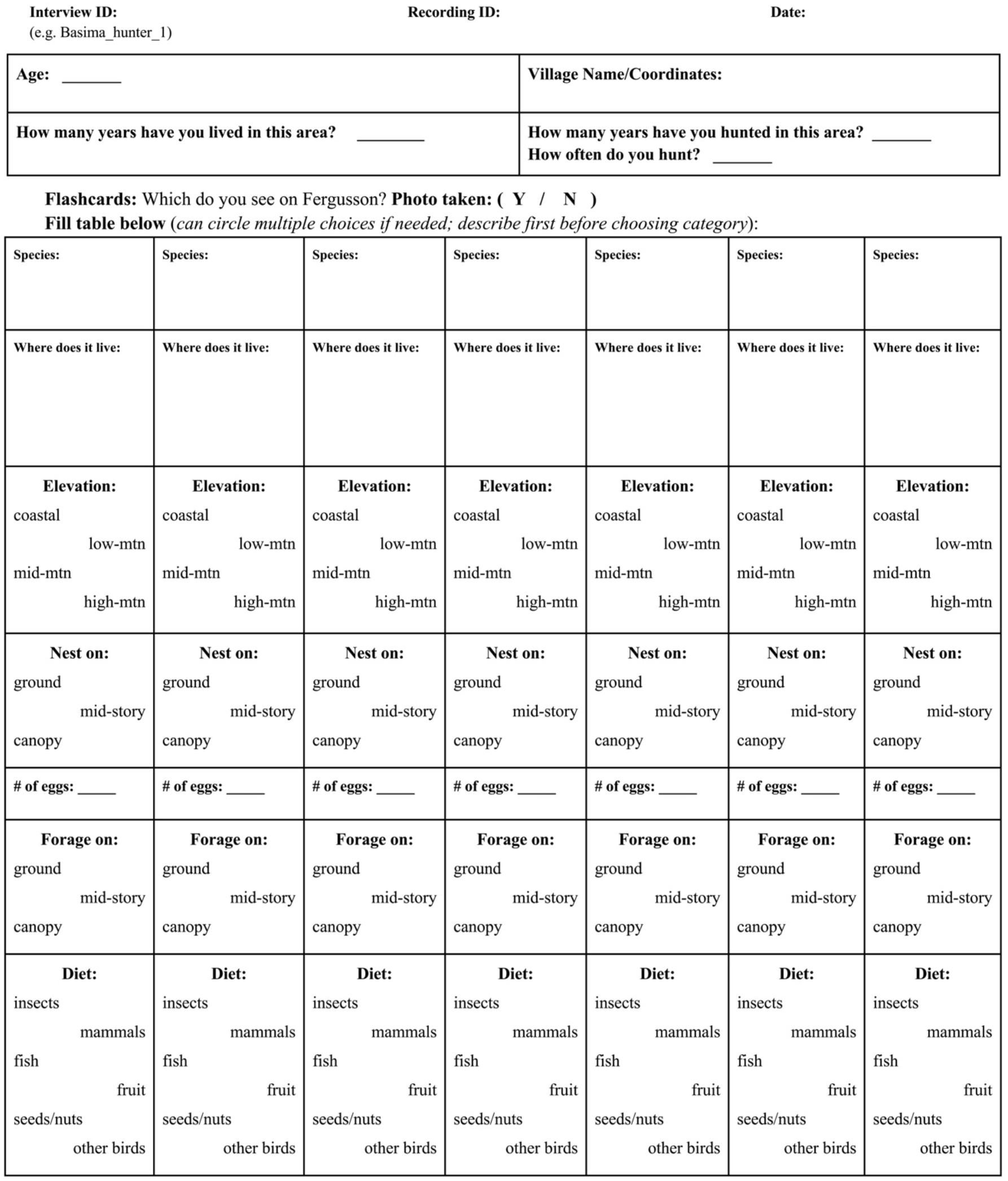

### Supplementary Material 1

Interview questionnaire, part 2.

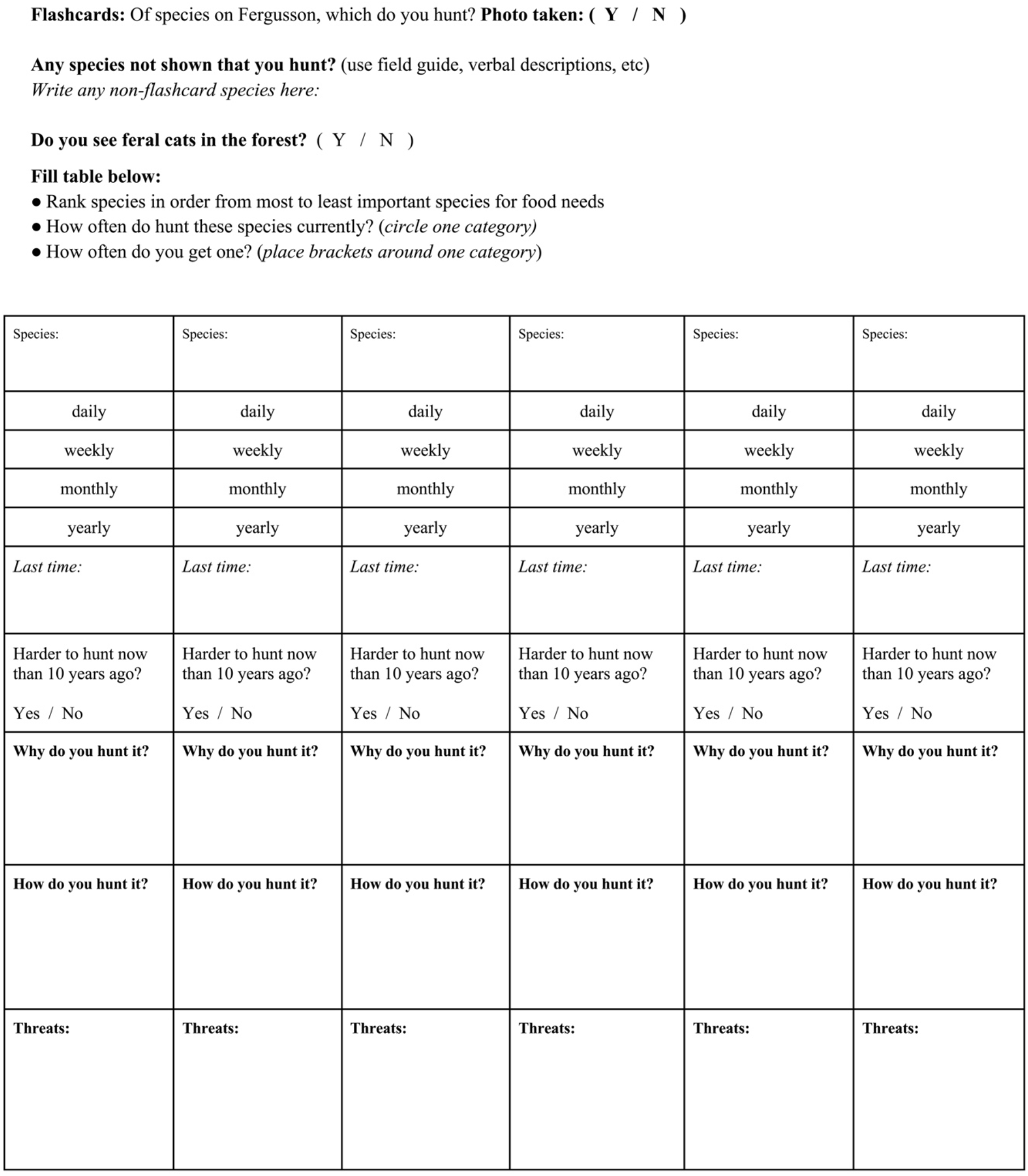

### Supplementary Material 2

Species table for visual and acoustic identification exercise.

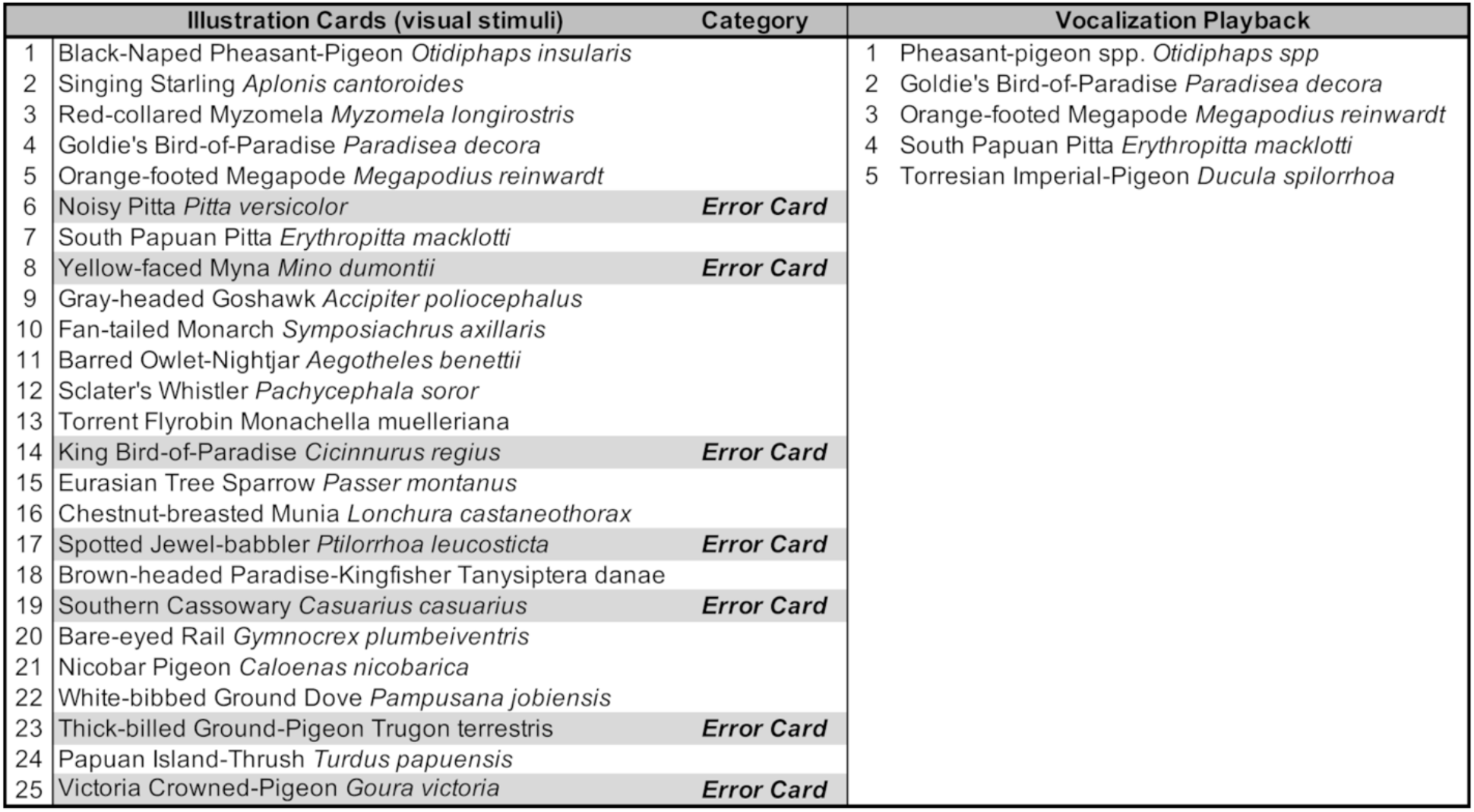

